# Comparison of the pathogenicity and virus shedding of SARS CoV-2 VOC 202012/01 and D614G variant in hamster model

**DOI:** 10.1101/2021.02.25.432136

**Authors:** Sreelekshmy Mohandas, Pragya D Yadav, Dimpal Nyayanit, Gururaj Deshpande, Anita Shete-Aich, Gajanan Sapkal, Sanjay Kumar, Rajlaxmi Jain, Manoj Kadam, Abhimanyu Kumar, Deepak Y Patil, Prasad Sarkale, Pranita Gawande, Priya Abraham

## Abstract

The emergence of SARS-CoV-2 variants has posed a serious challenge to public health system and vaccination programs across the globe. We have studied the pathogenicity and virus shedding pattern of the SARS-CoV-2 VOC 202012/01 and compared with D614G variant in Syrian hamsters. VOC 202012/01 could produce disease in hamsters characterized by body weight loss and respiratory tract tropism but mild lung pathology. Further, we also documented that neutralizing antibodies developed against VOC 202012/01 could equally neutralize D614G variant. Higher load of VOC 202012/01 in the nasal wash specimens was observed during the first week of infection outcompeting the D614G variant. The findings suggest increased fitness of VOC 202012/01 to the upper respiratory tract which could lead to higher transmission. Further investigations are needed to understand the transmissibility of new variants.

**One-Sentence Summary:** SARS-CoV-2 VOC 202012/01 infected hamsters demonstrated high viral RNA shedding through the nasal secretions and significant body weight loss with mild lung pathology compared to the D614G variant.

## Introduction

Severe acute respiratory syndrome coronavirus 2 (SARS-CoV-2) pandemic has significantly impacted the global healthcare and economy (*1*). Rapid spread of the virus, non-availability of any treatment or vaccines and insufficient healthcare facilities posed challenges in the beginning of the pandemic. Research on development of countermeasures against the disease was initiated in no time and now several vaccines are available across the globe like BNT162b2, mRNA-1273, ChAdOx1-S [recombinant], COVAXIN etc. (*2, 3*). The high mutation rate in RNA viruses helps them swiftly adapt to diverse environmental conditions posing a greater challenge in their prevention and control (*4, 5*). The continuous genomic surveillance of the circulating SARS-CoV-2 worldwide has given us clues about the genetic mutations acquired by the virus. The identification of such variants with potential of high transmission, disease severity, immunity evasion, resistance to therapeutic measures and ability to evade diagnostic tests is imperative to control the pandemic (*6*). Recently, various new SARS-CoV-2 variants have been reported worldwide which are of serious concern (*7, 8*).

SARS-CoV-2 is a single-stranded, positive-sense RNA virus that belongs to the genus *Betacoronavirus* of the *Coronaviridae* family. Its genome size is approximately 30 kbps, encoding different structural and non-structural proteins essential for the virus (*9*). The SARS-CoV-2 strains circulating during the early pandemics have been superseded with new variant strains (*7, 10*). Initially, SARS-CoV-2 strains with D614G mutation in the spike glycoprotein became predominant worldwide (*11*). This strain was associated with high viral loads in the upper respiratory tract than the earlier strains but didn’t produce severe disease (*12*). Recently, various viral strains of SARS-CoV-2 with mutations of concern like VOC 202012/01 (B.1.1.7 lineage) from the United Kingdom, VOC 202012/02 (B.1351 lineage) from South Africa, VOC 202101/02 (P.1) from Brazil, VOC 202102/02 i.e., a B.1.1.7 cluster with E484K mutation from the United Kingdom were reported (*7, 8*). Among this, VOC 202012/01 (B.1.1.7 lineage) from the United Kingdom defined by a range of 23 mutations, resulting in amino acid changes and three deletions in the virus has spread to 94 countries as of 16^th^ February 2021 with local transmission in at least 47 countries (*7, 13*). This variant has amino acid changes in the spike protein that included deletions (69/70,144Y) and replacements (N501Y, A570D, D614G, and P681H). The N501Y mutation in the Receptor Binding Domain (RBD) has been found to increase the binding affinity to Angiotensin Converting Enzyme 2 receptor by 10 times higher than that of the wild type RBD, which could increase the infectivity and transmissibility of the virus (*14*). Amino acid at position 501 which is in the RBD can possibly affect the efficacy of virus neutralization too as RBD accounts for 90% neutralizing activity against SARS-CoV-2 (*15*). The cumulative effect of all these mutations are still unknown and the initial reports from phylogenetic and transmission model analysis indicated that some of these mutations may influence the transmissibility of the virus in humans (*16*). The preliminary analysis data published by the New and Emerging Respiratory Virus Threats Advisory Group (NERVTAG) shows that there could be an increase in the risk of death associated with VOC 202012/01 variant compared to other non-VOC SARS-CoV-2 (*17*).

VOC 202012/02 reported from South Africa is characterized by spike protein mutations, K417N, E484K, N501Y, D614G and L242/A243/L244 deletions. Till date, the variant has been reported from 46 countries (*8, 18*). Another VOC was also reported from Brazil (20J/501Y.V3) belonging to P1 lineage causing the surge of infections (*10*). Recently, another VOC was reported from South West England called VOC 202102/02 i.e., a B.1.1.7 cluster with E484K mutation (*19*).

The continuous emergence of the SARS-CoV-2 variants is also becoming a serious threat to efficacy of the vaccines and monoclonal antibodies developed targeting spike protein which harbors mutations in the emerging variants. Investigations are ongoing to determine if these variants are associated with any changes in the severity of disease and transmission. This is the first study that investigated the pathogenesis of the SARS-CoV-2 strain VOC 202012/01 (B.1.1.7 lineage) in the golden (Syrian) hamster (*Mesocricetus auratus*) model and studied the pattern of the virus shedding of this new variant compared to the earlier SARS-CoV-2 D614G variant.

## Results

### Clinical observations in hamsters post infection with SARS-CoV-2 VOC-202012/01

Twenty Syrian hamsters were inoculated intranasally with 10^4.5^ TCID_50_ dose of SARS-CoV-2 VOC-202012/01 [‘hCoV-19/India/NIVP1 20203522/2020” (GISAID identifier: EPL_ISL_825088)] to determine its pathogenicity (Fig 1A). The infected hamsters demonstrated body weight loss with no apparent clinical signs (Fig.1B). Progressive body weight loss was observed till 7 days post-infection (DPI) (mean percent decrease = −7.34%) against a 2.7% mean increase in the control animals. Thereafter, a regaining trend in the body weight of hamsters from the infected group was observed till 14 DPI. The control animals (n=4) showed a gain of 6% till 14 DPI.

**Fig.1.**
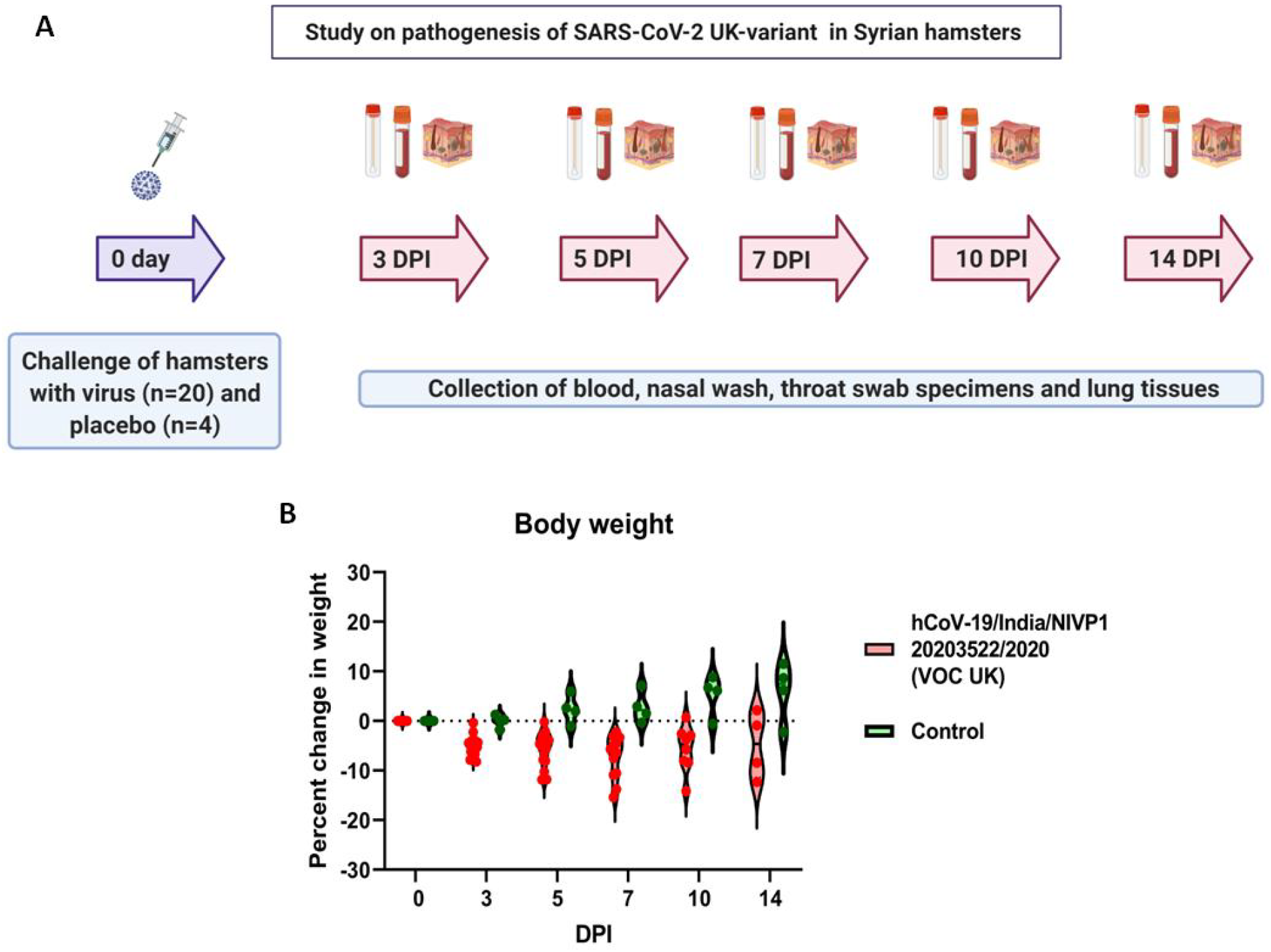
Study design and observations of the pathogenicity experiment of VOC 202012/21 in hamsters. (A) Study design of the pathogenicity of VOC 202012/21 in Syrian hamsters (B) The percent body weight difference observed in animals post VOC 202012/01 challenge and in mock control till 14 days.

### Immune response in hamsters post infection with SARS-CoV-2 VOC-202012/01

The infected animals exhibited anti-SARS-CoV-2 IgG response from 7^th^ DPI which persisted till day 14 (Fig 2A-C). The response was found predominantly of IgG2 type. The serum samples of the infected hamsters were checked for their neutralization ability against both homologous (VOC-202012/01) and heterologous (D614G variant) SARS-CoV-2 strains. Although on 5 DPI, the neutralization titer differed significantly with a higher titer against the VOC 202012/01, the samples showed comparable neutralization titer from 7 DPI till 14 DPI with both the variants (Fig 2D).

**Fig.2.**
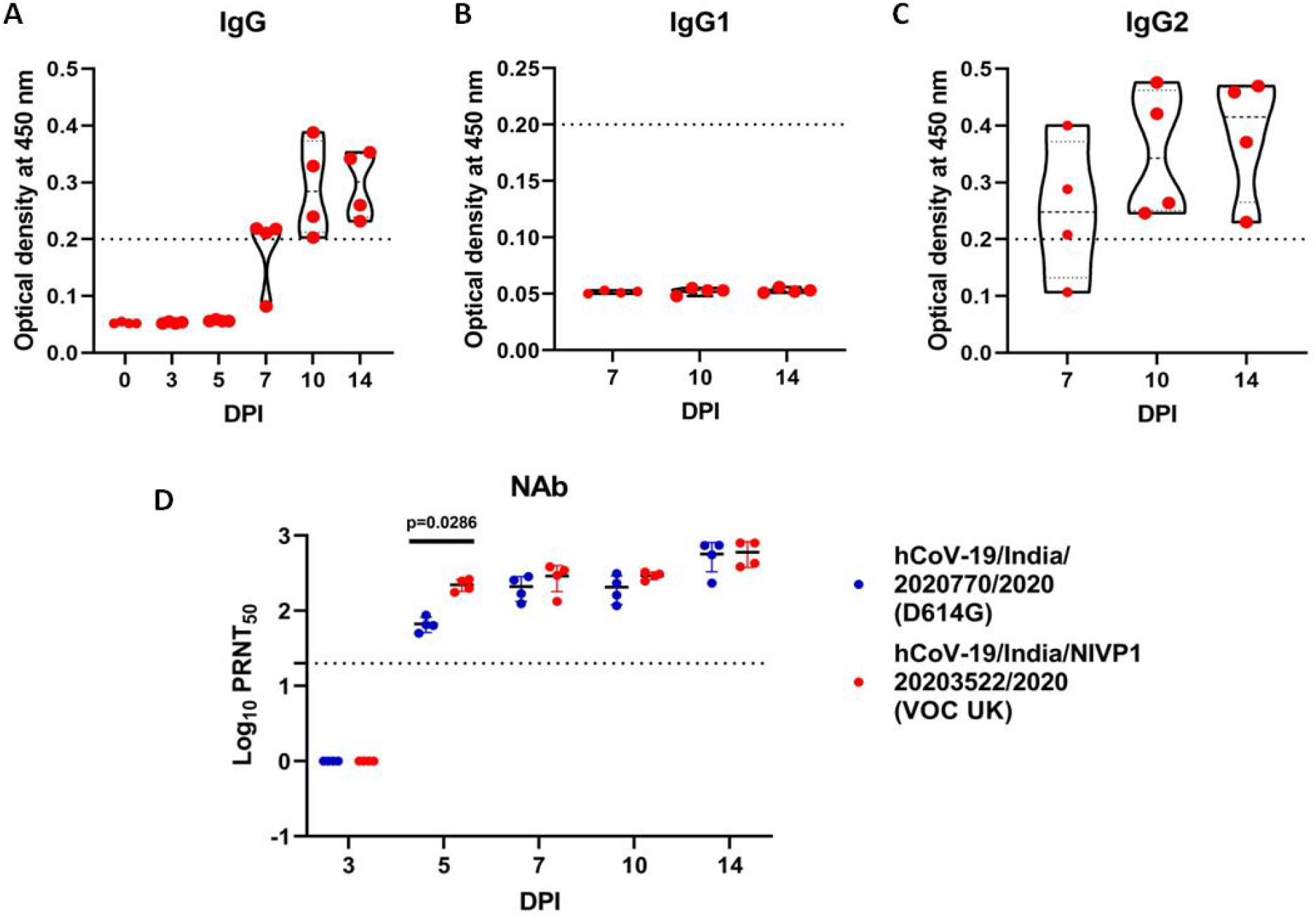
Humoral immune response developed in hamsters post infection with VOC 202012/01. (A) Anti-SARS-CoV-2 IgG responses in hamsters post infection. (B) IgG1 and (C) IgG2 response in serum samples post infection in hamsters at 7, 10 and 14 DPI. (D) Neutralizing antibody titers of hamster serum samples against SARS-CoV-2 VOC 202012/01 and D614G variant. The dotted line indicates the limit of detection of the assay.

### SARS-CoV-2 viral load in hamsters post infection with SARS-CoV-2 VOC-202012/01

The respiratory tract tropism of the virus was evident with the high viral genomic RNA (gRNA) load in the upper and lower respiratory tract (Fig 3A-E). Among these, lungs (1.7 x 10^10^/ml) showed highest mean viral load followed by nasal wash (5.4 x 10^8^/ml), nasal turbinates (2.6 x 10^8^/ml), trachea (2.3 x 10^8^/ml) and throat swabs (2.9 x 10^7^/ml) on 3 DPI. A decreasing trend was observed in the gRNA load in the nasal wash, throat swab and trachea from 3 to 14 DPI. The gRNA peak in the lungs and nasal turbinate specimens were observed on 5^th^ DPI which declined further till 14 DPI. The gRNA load reached up to 8.8 x10^6^/ml, 3.7 x 10^6^/ml and 4.3 x 10^4^/ml in lungs, nasal turbinates and nasal wash respectively on 14 DPI. Lungs showed an average viral titer of 10^4.2^ and 10^2.6^ TCID50/ml on 3 and 5 DPI respectively, whereas nasal turbinates showed an average of 10^2.6^, 10^2.5^ and 10^2^ TCID50/ml on 3, 5 and 7 DPI. No titer could be observed in these samples at 10 and 14 DPI. Lungs and trachea samples of all hamsters were found positive for subgenomic RNA (sgRNA) on 3 DPI whereas only two out of four hamsters showed sgRNA positivity in nasal wash and turbinate on 3 DPI (Fig 3F-J). Lungs samples were found to be consistently positive for sgRNA on 5^th^ DPI too but only one hamster was positive for the sgRNA in the nasal wash and nasal turbinate. Other organs like small intestine, kidney, liver and brain showed gRNA positivity from some animals on 3 and 5 DPI. However, sgRNA could not be detected in any of these organs.

**Fig.3:**
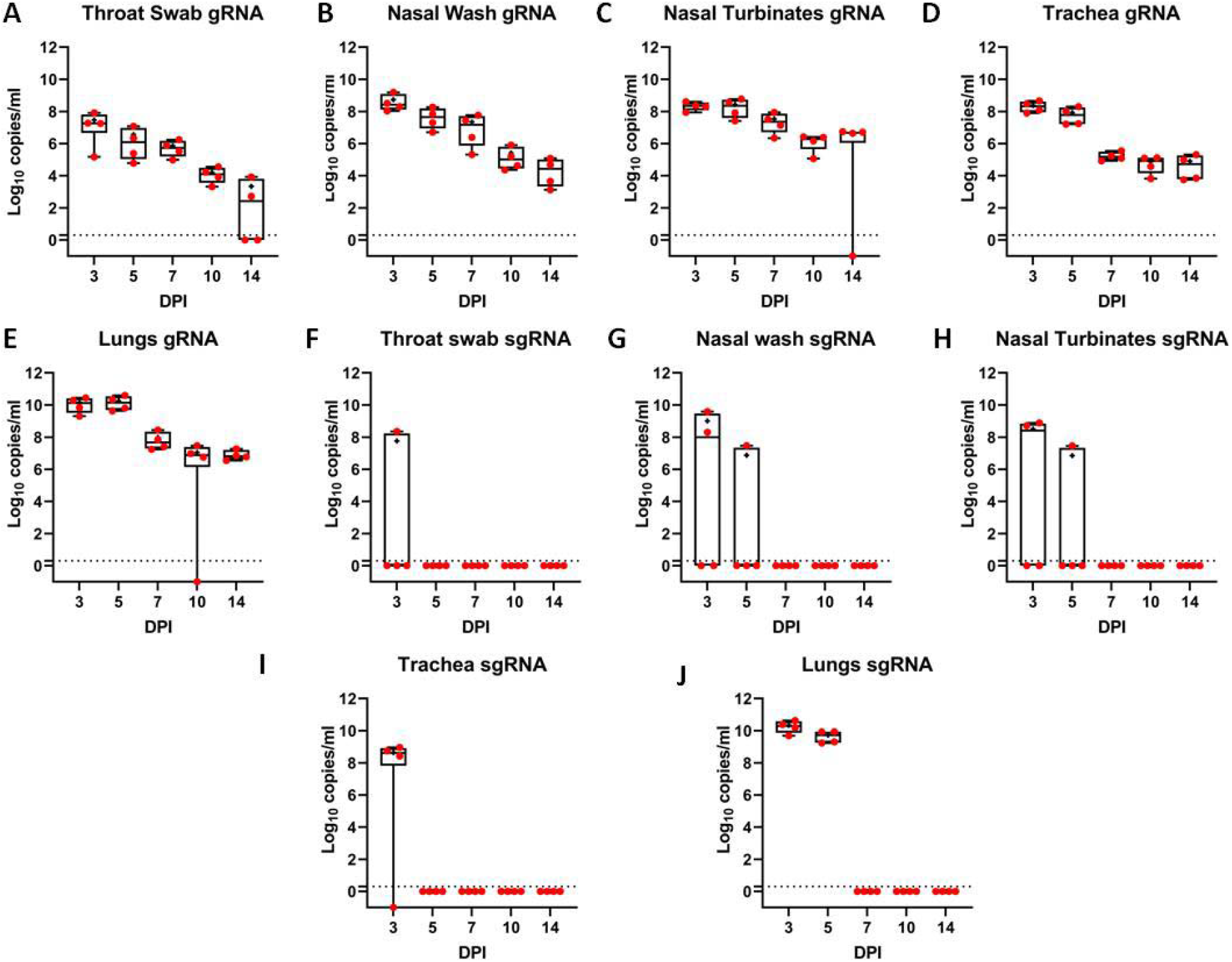
SARS-CoV-2 genomic and subgenomic RNA loads tested by E gene real time RT-PCR in hamsters infected with SARS-CoV-2 VOC 202012/01. The viral gRNA load in throat swab (A), nasal wash (B), nasal turbinates (C), trachea (D) and lungs (E) samples collected sequentially on 3, 5, 7, 10 and 14 DPI from VOC 202012/01 infected hamsters represented as mean ± standard deviation (SD) of copy numbers/ml. The viral sgRNA load in throat swab (F), nasal wash (G), nasal turbinates (H), trachea (I) and lungs (J) samples collected sequentially on 3, 5, 7, 10 and 14 DPI from VOC-202012/01 infected hamsters represented as mean ± standard deviation (SD) of copy numbers/ml.

### Pathological changes in lungs post infection with SARS-CoV-2 VOC 202012/01

On gross examination of organs during necropsy, lungs showed congestive changes and hemorrhagic foci of multiple sizes from 5 DPI which persisted till 10 DPI (Fig 4A). The gross changes in the lungs appeared resolved on 14 DPI. Histopathologically, lungs showed foci of consolidation and mild exudative changes on 5 DPI which progressed to mild alveolar septal thickening characterized by hyperplasia and inflammatory cell infiltration and consolidation on 7 DPI. Lungs sections of the 10 and 14 DPI showed only focal areas of mononuclear cell infiltration and consolidation in the peribronchial region without any alveolar damage (Fig. 4B-D). Our earlier study with 1 x 10^4.5^ TCID50 of SARS-CoV-2 D614G variant produced more severe pneumonic changes in lungs (Table S1) (*20*). Congestion and hemorrhages were observed as early as 3 DPI in D614G variant infected animals and more pronounced alveolar septal thickening, consolidation and inflammatory infiltration were also observed (4E, F).

**Fig.4:**
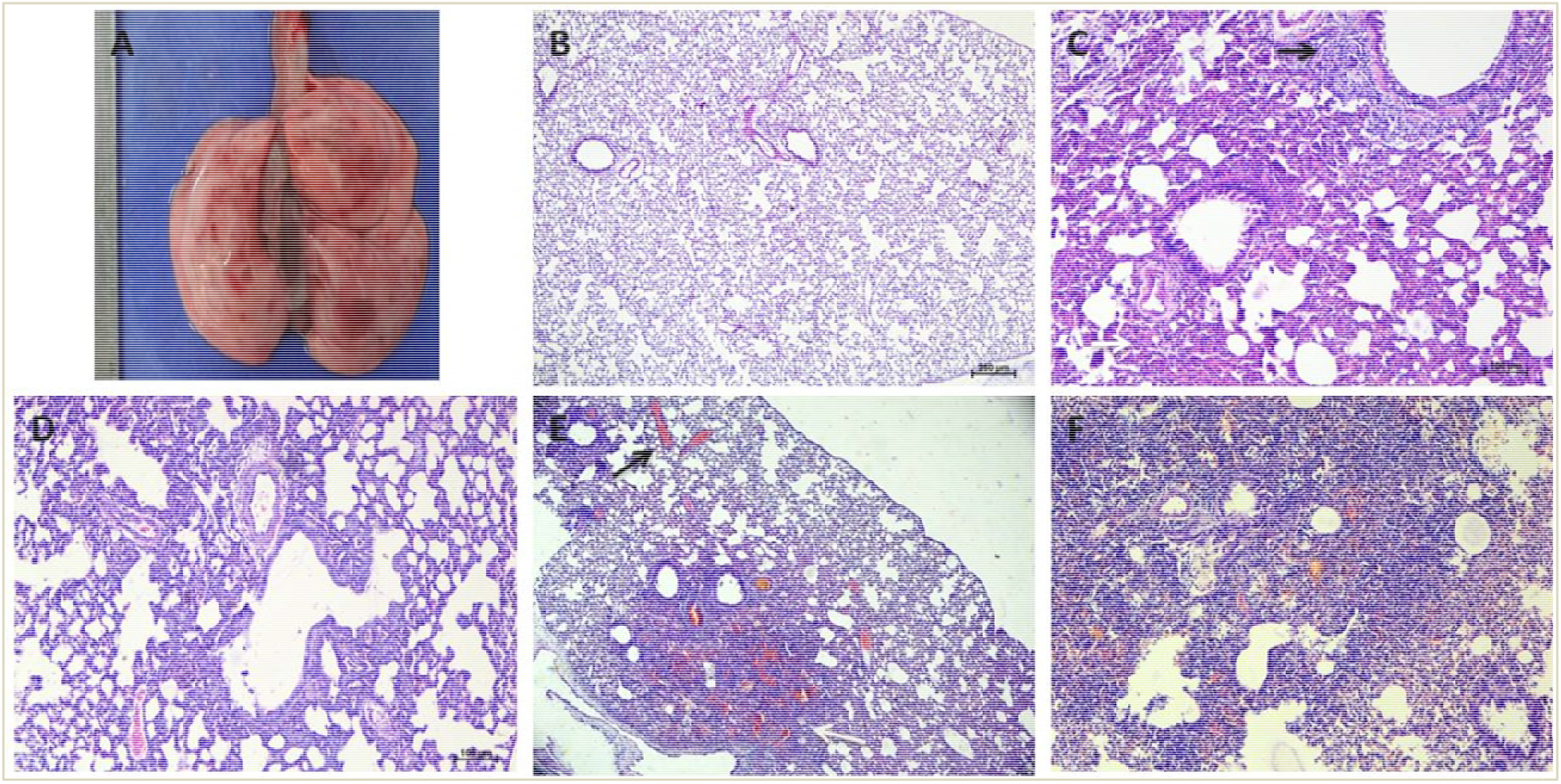
Pathological changes in lungs of hamsters’ post-SARS-CoV-2 VOC 202012/01 and D614G variant infection. (A) Lungs of hamster infected with SARS-CoV-2 VOC 202012/01 on 7 DPI showing congestion and multiple hemorrhagic foci. (B) Lungs of hamster infected with SARS-CoV-2 VOC 202012/01 showing normal morphological features on 3 DPI, scale bar=250μm. (C) Lungs of hamster infected with SARS-CoV-2 VOC 202012/01 showing peribronchiolar mononuclear cell infiltration (black arrow) and consolidative changes in the alveolar parenchyma (white arrow) on 10 DPI infected with SARS-CoV-2 VOC 202012/01, scale bar=100μm. (D) Lungs of the hamster infected with SARS-CoV-2 D614G showing normal alveolar parenchyma and mild peribronchial consolidation on 14 DPI, scale bar=100μm. (E) Lungs showing severe congestion and a focus of consolidation and alveolar damage on 3 DPI with SARS CoV-2 D614G variant, scale bar=250μm. (F) Lungs showing diffuse congestive changes, hemorrhages alveolar septal thickening, and mononuclear infiltration on 14 DPI with SARS CoV-2 D614G variant scale bar=250μm.

### Serum cytokine profile in hamsters post infection with SARS-CoV-2 VOC-202012/01

We have assessed the serum cytokine levels in the infected hamsters on 3, 7 and 14 DPI and compared with SARS-CoV-2 D614G variant from an earlier study where hamsters were intranasally infected with 10^5.5^ TCID50/ml of the virus. But no significant difference could be observed in the cytokine levels in hamsters infected with both variants (Fig.S1).

### Comparison of virus shedding in hamsters infected with VOC-202012/01 and D614G variant

We inoculated six hamsters each intranasally with 10^4.5^ TCID50 doses of the SARS-CoV-2 VOC 202012/01 and SARS-CoV-2 D614G variant to determine the virus shedding pattern of both the variants through the nasal tract, throat and faeces (Fig.5A). The infected hamsters were observed for 14 days. Significant body weight loss was observed in hamsters infected with VOC-202012/01 compared to the D614G variant (Fig.5B). Persistent viral gRNA detection was noted in the throat, nasal wash and faecal samples of these animals in a decreasing trend till 14 DPI (Fig.4B-D). Nasal wash samples showed high viral gRNA load compared to the throat or fecal samples in hamsters infected with both strains. Subgenomic RNA could be detected in these samples only on 2 and 4 DPI (Fig 4E-F). The gRNA loads in the nasal wash samples were significantly high on 4, 6 and 8 DPI (Fig. 4C). Virus titration performed on throat swab and nasal wash samples till 10 DPI demonstrated the persistent virus titer and no statistical significance in virus titer was observed among the two variants (Fig. S2).

**Fig.5:**
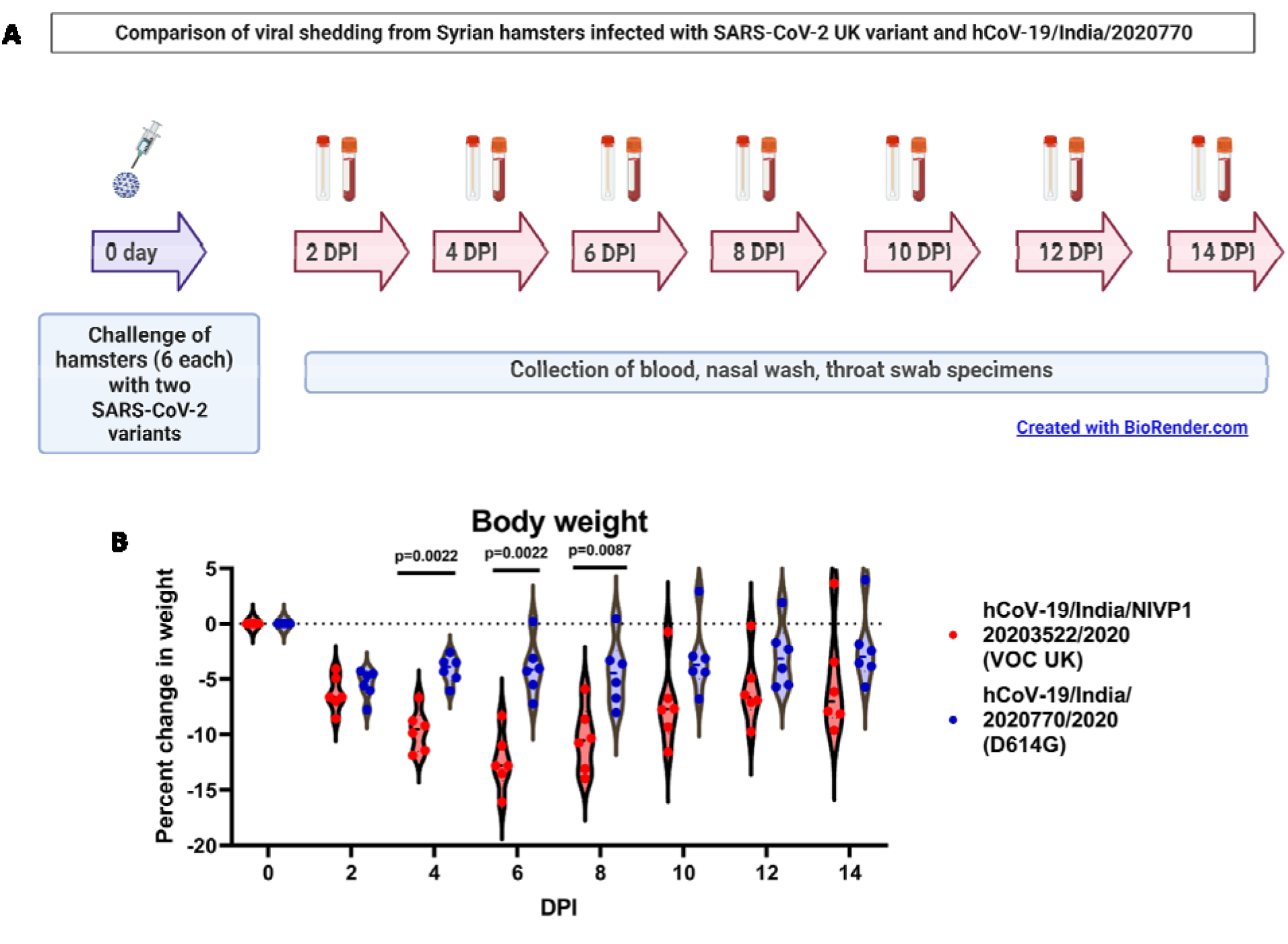
Study design and observations of the virus shedding comparison study in hamsters infected with SARS-CoV-2 VOC 202012/01 and D614G variant. (A) Study design (B) Percent bodyweight change in hamsters infected (mean ± standard deviation). The statistical significance was assessed using the non-parametric Mann-Whitney test between the two groups; p-values less than 0.05 were considered to be statistically significant.

**Fig.6:**
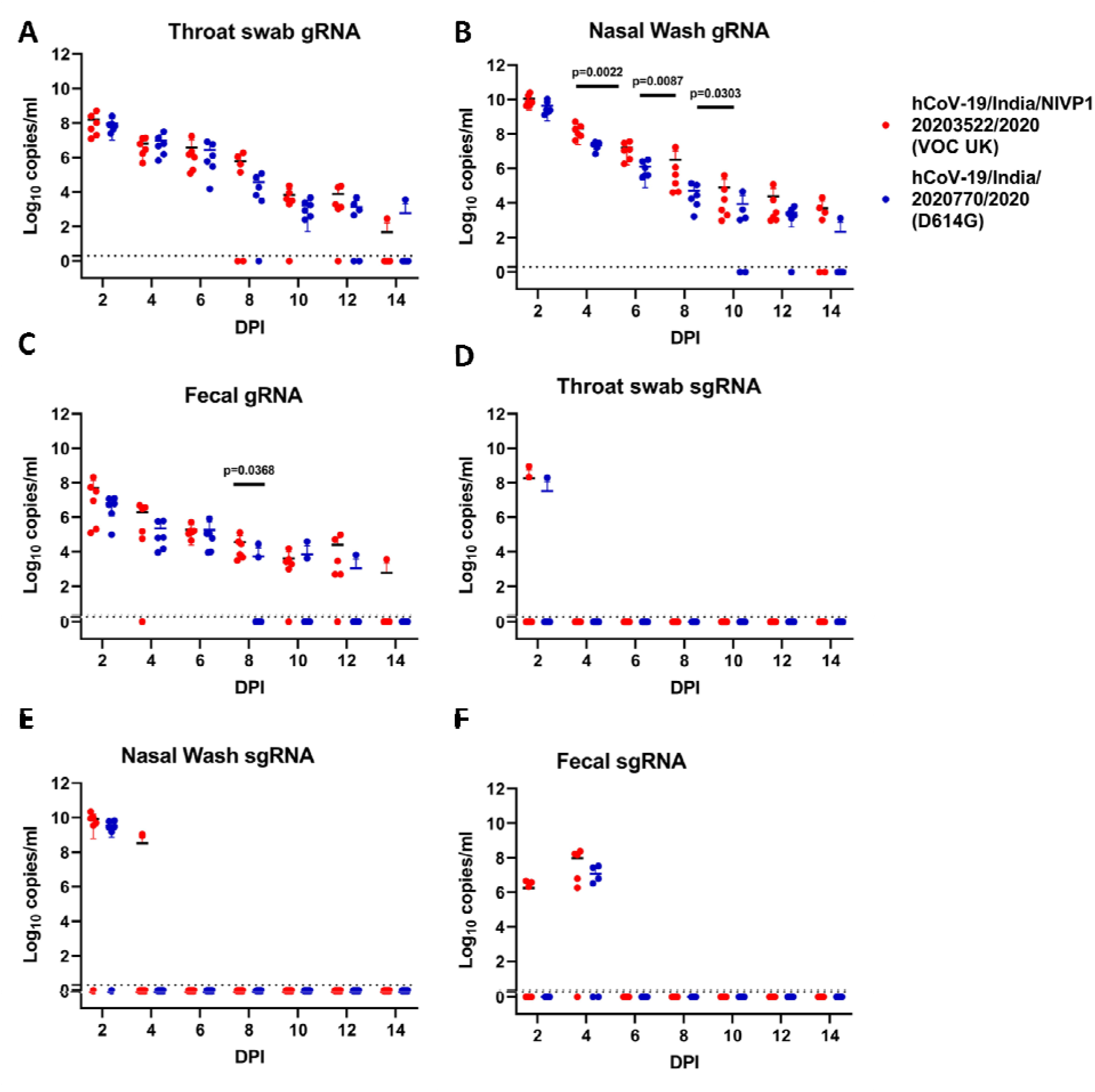
SARS-CoV-2 genomic and subgenomic RNA loads in hamsters infected with SARS-CoV-2 VOC 202012/01 and D614G variant. (A) The viral gRNA load in throat swab (B) nasal wash and (C) fecal samples collected sequentially on 2, 4, 6, 8, 10, 12 and 14 DPI from VOC-202012/01 and D614G infected hamsters represented as mean ± standard deviation of copy numbers/ml). (D) The viral sgRNA load in throat swab (E) nasal wash and (F) fecal samples collected sequentially on 2, 4, 6, 8, 10, 12 and 14 DPI from VOC-202012/01 and D614G infected hamsters represented as mean ± standard deviation of copy numbers/ml. The statistical significance was assessed using the non-parametric Mann-Whitney test between the two groups; p-values less than 0.05 were considered to be statistically significant.

## Discussion

Considering the worldwide concern and speculation of the high transmission and disease severity related to VOC20212/01, we assessed the disease severity and virus shedding pattern of VOC 20212/01 in hamsters. Persistent body weight loss was observed in hamsters as reported with other variants of SARS-CoV-2 (*21-24*). We could observe a significant higher percent body weight loss in hamsters infected with same titer of VOC 202012/01 compared to D614G variant indicating its pathogenicity. The comparable serum neutralization titer of VOC 202012/01 infected hamsters against heterologous virus strain is in line with the available evidence of VOC 202012/01 infected humans suggesting no strong association with escape from the natural or vaccine acquired immunity (*6, 25, 26*). The IgG response was predominantly IgG2 type as observed earlier in hamsters infected with SARS-CoV-2 D614G variant and in ß-propionolactone inactivated SARS-CoV-2 (BBV152) vaccinated hamsters (*20*).

In our earlier study with SARS-CoV-2 D614G variant in hamsters, the peak gRNA load in lungs and nasal turbinate was detected on 3 DPI and further decrease on subsequent days (*22*). Here, we have observed gRNA peak in the lungs and nasal turbinate specimens on 5 DPI, indicating a delay in virus clearance compared to the earlier strain. The virus clearance trend appeared proportional to the increasing neutralizing antibody titer from 7 DPI. We have observed virus titer in lungs and nasal turbinate only on 3 and 5 DPI in our earlier study with the D614G variant with a higher TCID50 dose of infection (*22*). Here nasal turbinates showed presence of live virus till 7 DPI indicating a longer persistence of the virus which could probably increase the window period of transmission. sgRNA could not be detected in any of the organs other than the respiratory tract samples substantiating the respiratory tissue tropism of the strain similar to the earlier studied strains of SARS CoV-2. Although, we observed higher viral gRNA load in the respiratory tract samples, only mild histopathological changes were observed in the lungs with an inoculum of 1 x 10^4.5^ TCID_50_ dose. The severity of disease reported in hamsters by SARS-CoV-2 varied with the dose of virus inoculums (*21–24*). Earlier studies have shown that even with lower dose of virus inoculums, comparable viral RNA load was observed in 3 to 4 DPI as that of higher dose but the lung pathological and body weight changes vary with increasing dose suggesting a dose dependant disease severity in hamsters (*20, 21, 24, 27*). Even though we could observe an increased percent body weight loss indicating the severity but lung histopathological changes were not severe. This finding is in contrary to the recent reports by NERVTAG which showed the evidences from analysis of multiple different datasets that infection with VOC 202012/01 is associated with an increased risk of hospitalization and death compared to infection with non-VOC, even though there are potential limitations in the datasets used which includes the representativeness, potential biases in case enrolment, confounders etc (*17*). The amount of virus shedding through the body secretions and excretions mainly contributes to the virus transmission. The high gRNA was observed in the nasal wash samples consistently during the first week post infection in VOC 202012/01 hamsters although comparable virus titers were observed with both variants in these samples.

In conclusion, the intranasal infection with the SARS-CoV-2 VOC 202012/01 produced disease in hamsters characterized by body weight loss, infection of the upper and lower respiratory tract and mild lung pathology. No increased disease severity in terms of lung pathology could be observed in hamsters as speculated in humans with VOC 202012/01 but significant decrease in body weight was observed. There was no difference in the neutralization potential against D614G variant. The higher viral RNA load in nasal washes of the SARS-CoV-2 VOC-202012/01 infected hamsters could be a supporting evidence for the increased transmissibility reported with this strain. Further, transmissibility studies need to be explored.

## Supplementary Materials

Materials and Methods

Figs. S1 to S2

Table S1

## Materials and Methods

### Virus strains

“hCoV-19/India/NIVP1 20203522/2020” (GISAID identifier: EPL_ISL_825088) with the characteristic amino acids changes similar to the 20I/501Y.V1 (GR clade), also called as Variant of Concern (VOC) 202012/01 with a tissue culture infective dose 50 (TCID50) of 10^5.5^/ ml and “hCoV-19/India/2020770/2020 (GISAID identifier: EPL_ISL_825088), the SARS-CoV-2 NIV 2020-770 strain (G clade) possessing D614G mutation with a tissue culture infective dose 50 (TCID50) of 10^6.5^/ ml were used in the study. Both the strains were passaged up to three times in Vero CCL-81 cells before use. The strains were isolated from nasal swab or throat swab samples of COVID-19 patients at Indian Council of Medical Research (ICMR)-National Institute of Virology (NIV), Pune (*28, 29*).

### Ethics statement

The study was approved by the Institutional Animal Ethics Committee and Institutional Biosafety Committee of ICMR-NIV, Pune with the approval no. of NIV/IAEC/2021/MCL/01 and NIVIBSC/05.01.2021/02 respectively. All the experiments were performed as per the guidelines of the Committee for the Purpose of Control and Supervision of Experiments in Animals, Government of India (*30*).

### Experiments in Syrian hamsters

Twenty-four (12 male and 12 female) Syrian hamsters of 8-10-week-old age were used for the pathogenicity experiment. The hamsters were housed in individually ventilated cages with ad libitum food and water. Twenty hamsters were challenged with 0.1 ml of 10^5.5^ TCID50/ml hCoV-19/India/NIVP1 20203522/2020 (VOC 202012/01) intranasally in the containment facility of ICMR-NIV, Pune under isoflurane anesthesia. Four hamsters were kept as mock control and were inoculated with sterile tissue culture media. Animals were monitored daily for any clinical signs and the body weights were monitored till 14 days. Four hamsters from each group were euthanized on 3, 5, 7, 10 and 14 DPI to collect blood, organ and swab samples for viral RNA estimation, titration, histopathology, and immunological analysis.

For the comparison study of virus shedding in hamsters, twelve (6 male and 6 female) Syrian hamsters of 8-10-week-old age were used in the study. The hamsters were housed in individually ventilated cages with ad libitum food and water. Six hamsters (3 males and 3 females) each were challenged with 0.1 ml of 10^5.5^ TCID50/ml SARS-CoV-2 VOC-202012/01 and D614G variant intranasally in the containment facility under isoflurane anesthesia. The animals were monitored for any clinical signs and the body weight loss. Throat swabs, nasal wash and facecal samples were collected in 1 ml virus transport media on every alternate DPI till 14 days for viral load estimation.

### Real-time Reverse Transcriptase-Polymerase Chain Reaction (rRT-PCR)

Nasal wash, throat swab and faecal samples were collected from hamsters in 1ml viral transport medium. Organ samples were weighed and triturated in 1 ml sterile tissue culture media using a tissue homogenizer (Eppendorf, Germany). Two hundred microliters of the tissue homogenate/ swab specimens were used for RNA extraction using MagMAX™ Viral/Pathogen Nucleic Acid Isolation Kit as per the manufacturer’s instructions. Real-time RT-PCR was performed for E gene of SARS-CoV-2 using the primers, Forward: ACAGGTACGTTAATAGTTAATAGCGT, Reverse: ATATTGCAGCAGTACGCACACA and probe: FAM-ACACTAGCCATCCTTACTGCGCTTCG-BBQ (*31*). The cycling conditions used were 55°C for 30 minutes for reverse transcription, followed by 95°C for 3 minutes and 45 cycles of 95°C for 15 s, 58°C for 30 s. For detection of sgRNA of E gene published primers (Forward: CGATCTCTTGTAGATCTGTTCTCT-3, Reverse: ATATTGCAGCAGTACGCACACA-3 and probe: FAM CACTAGCCATCCTTACTGCGCTTCG-3) were used (*32*). The cycling conditions used were 50°C for 30 minutes at for reverse transcription, followed by 95°C for 2 minutes and 45 cycles of 95°C at 15 seconds, 50°C for 1minute.

### Virus titration

The lungs, nasal turbinate, throat swab and nasal wash samples were used for virus titration in Vero CCL-81 cells. Tissue homogenates were prepared using a tissue homogenizer (Eppendorf, Germany) in sterile Minimum Essential Media (Gibco, USA) followed by centrifuging at 1984 g for 10 minutes. The supernatant is further used for the titration. Hundred micro liters (μl) of the tissue homogenate/ swab specimens were added onto a 24-well plate with Vero CCL81 monolayers and incubated at 37°C for one hour. The cell monolayer was washed with phosphate buffered saline (PBS) post incubation after the removal of the media. The plates were subsequently incubated with maintenance media containing 2% fetal bovine serum (FBS) in a CO2 incubator at 37°C. The plates were examined daily for any cytopathic effects (CPE) using an inverted microscope (Nikon, Eclipse Ti, Japan). The supernatant from the wells showing CPE was harvested and confirmed by real-time RT-PCR. The titers were calculated using the Reed and Muench method and were expressed at TCID50/ml (*33*).

### Enzyme-linked Immunosorbent Assay

The serum samples collected on day 3, 5, 7, 10 and 14 DPI were tested for IgG antibodies by hamster anti-SARS-CoV-2 IgG ELISA. The assay procedure in brief is as follows. Ninety-six well microtiter plates were coated with inactivated SARS-CoV-2 antigen in carbonate buffer of pH 9.5 overnight at 4 °C. The wells were blocked with liquid plate sealer for two hours at room temperature (25-30°C) and further washed with phosphate-buffered saline with 0.05% Tween 20 (PBS-T). The plates were incubated at 37°C for one hour with 100μl of diluted hamster serum samples (1: 100). After 5 washes with PBS-T, anti-hamster IgG-horse radish peroxidase1:3000 (Thermoscientific, USA) were added and incubated for 1 hour at 37°C. The plate was washed with PBS-T for five times. Hundred μl of substrate, 3’,3’5,5’-tetramethylbenzidine (TMB) was added to each well following the wash. Absorbance was measured at 450 nm following the termination of color developed. Optical density of more than 0.2 and positive/negative (P/N) ratio above 1.5 was considered positive. For antibody sub-typing, the same protocol was followed except that of the use of biotinylated anti-Syrian hamster IgG1 /IgG2 antibodies in 1:10000 dilution (BD biosciences, USA) and Streptavidin-horseradish peroxidase 1:8000 (Thermo-scientific, USA) for detection.

### Plaque Reduction Neutralization test (PRNT)

The assay was performed using hamster serum samples collected on 3, 5, 7, 10 and 14 DPI with both strains of virus used in the study i.e., SARS-CoV-2 VOC-202012/01 and SARS-CoV-2 D614G variant (NIV 2020-770 strain). The serum samples were heat inactivated at 56°C for 30 minutes in a water bath. Four-fold serial dilution of hamster serum samples were mixed with an equal amount of virus containing a 50-60 plaque forming units/0.1 ml and were incubated at 37°C for 1 hour. 0.1 ml of this mixture was added in a 24-well tissue culture plate containing a confluent monolayer of Vero CCL-81 cells with intermittent shaking. The plates were incubated at 37°C for 60 minutes and the mixture was aspirated. The culture overlay medium containing 2% carboxymethyl cellulose with 2% FBS in 2X minimal essential media was added onto the cells. The plates were incubated at 37°C CO2 incubator for 4 days. The overlay medium was decanted and the cells were stained by 1% amido black. The plaque number was counted and PRNT50 titers were calculated using a log probit regression analysis.

### Histopathology

Lungs samples collected during necropsy were fixed in 10% neutral buffered formalin. The tissues were processed using an automated tissue processor. The processed tissues were embedded in paraffin and were sectioned using an automated microtome (Leica, Germany) and were stained by hematoxylin and eosin staining.

### Serum cytokine analysis

The serum cytokine levels (IL-2, IL-4, IL-6, IL-10, IL-17A, TNF-α and IL-12) were assessed in hamsters serum samples collected at 3, 7 and 14 DPI. A commercial ELISA (Immunotag, USA) was used for the hamster specific cytokine quantitation. For this, plates precoated with hamster specific cytokine antibody were used and a streptavidin based HRP system was used for detection. The absorbance was measured at 450 nm.

### Data analysis

For analysis of the data, Graph pad Prism version 8.4.3 software was used. For statistical analysis; non parametric Mann Whitney tests were used and the p-values less than 0.05 were considered to be statistically significant.

---

**Fig S1:**
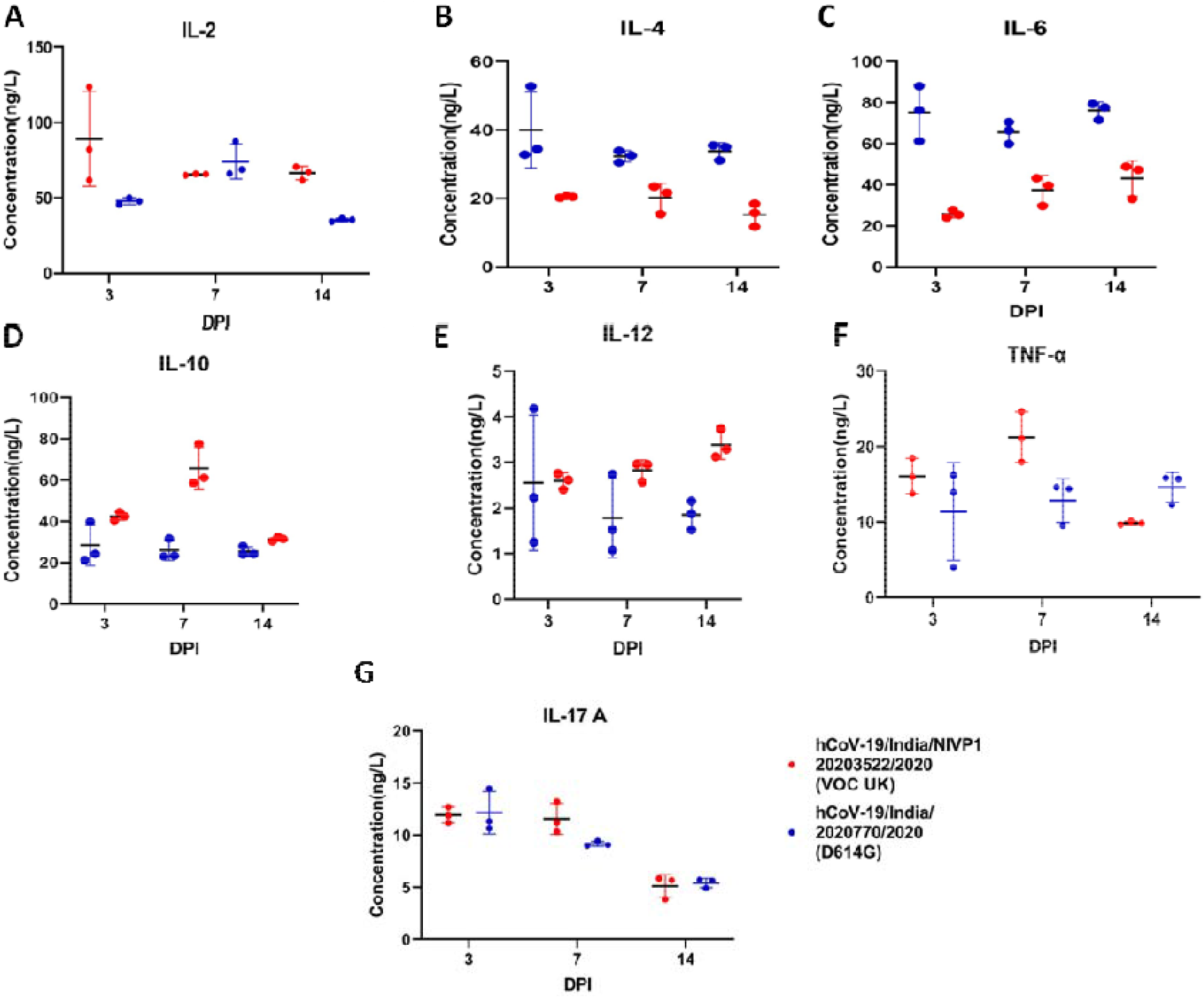
Comparison of serum cytokine levels in hamsters post infection. (A) IL-2 (B) IL-4 (C) IL-6 (D) IL-10 (E) IL-12 (F) TNF-α (G) Il-17A in hamsters on 3, 7 and 14 DPI with SARS-CoV-2 VOC 202012/01 and D614G variant. For statistical analysis; non parametric Mann Whitney tests were used and the p-values less than 0.05 were considered to be statistically significant.

**Fig S2:**
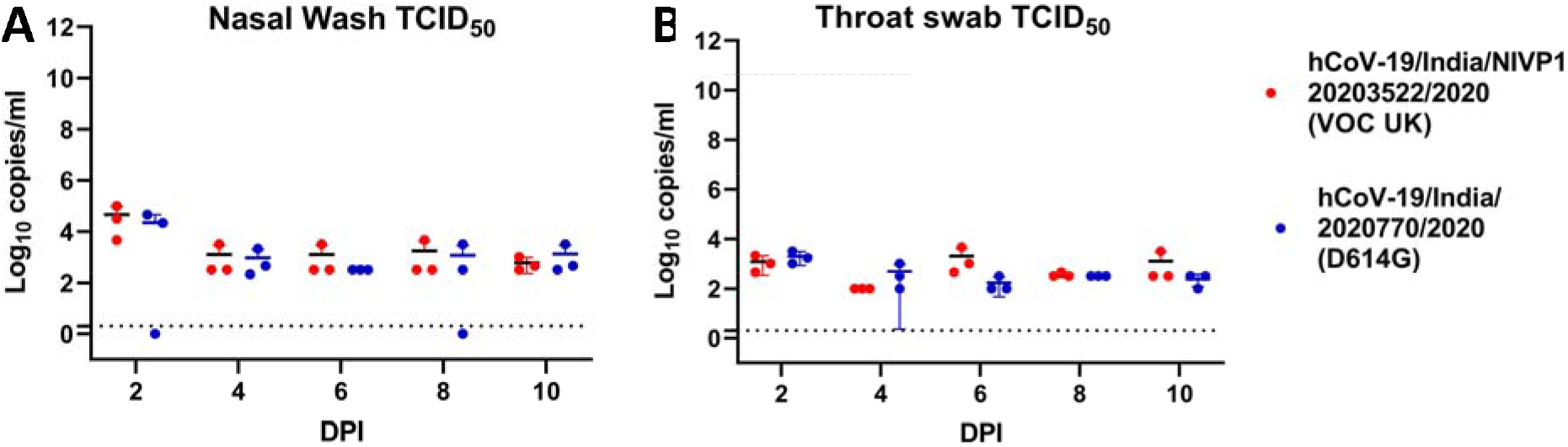
Virus titration of the nasal wash and throat swab samples collected sequentially on 2, 4, 6, 8, and 10 DPI from VOC 202012/01 and D614G variant infected hamsters. (A) Virus titer in the nasal wash samples expressed as TCID50/ml plotted in a scatter plot along with mean ± standard deviation. (B) Virus titer in the throat swab samples expressed as TCID50/ml plotted in a scatter plot along with mean ± standard deviation. For statistical analysis; non parametric Mann Whitney tests were used and the p-values less than 0.05 were considered to be statistically significant.

**Table S1:** The histopathological criteria and scoring of the lung samples collected post infection from hamsters. The severe changes were graded as +4, moderate as +3, mild as +2, minimal as +1 and no lesions as ‘0’.

| Virus used | DPI | Histopathological criteria and scoring of lungs |  |  |  |  |
| --- | --- | --- | --- | --- | --- | --- |
|  |  | Congestion and hemorrhages | Degenerative & necrotic changes | Consolidation, thickening of alveolar septa | Edematous changes | Cellular infiltration |
| SARS-CoV-2 D614G variant | 3 | +2 | 0 | +2 | +1 | +2 |
|  |  | +1 | 0 | +1 | 0 | +1 |
|  |  | +3 | 0 | +2 | +1 | +2 |
|  | 14 | +1 | +1 | +2 | 0 | +2 |
|  |  | +1 | +1 | +1 | 0 | +2 |
|  |  | +1 | +1 | +1 | +1 | +2 |
| SARS-CoV-2 202012/01 | 3 | 0 | 0 | 0 | +1 | 0 |
|  |  | 0 | 0 | 0 | 0 | 0 |
|  |  | 0 | 0 | 0 | 0 | 0 |
|  | 5 | 0 | 0 | +1 | +1 | +1 |
|  |  | +1 | 0 | +1 | +1 | +1 |
|  |  | +1 | 0 | +1 | +1 | +1 |
|  | 7 | 0 | 0 | +2 | +1 | +2 |
|  |  | 0 | 0 | +1 | 0 | +1 |
|  |  | 0 | 0 | +2 | +1 | +1 |
|  | 10 | 0 | 0 | +1 | 0 | 0 |
|  |  | 0 | 0 | +1 | 0 | +1 |
|  |  | 0 | 0 | +1 | 0 | +1 |
|  | 14 | 0 | 0 | 0 | 0 | 0 |
|  |  | 0 | 0 | +1 | 0 | +1 |
|  |  | 0 | 0 | +1 | 0 | +1 |

## Acknowledgments

The authors acknowledge the support received from Dr Chandrasekhar Mote, Assistant Professor, Department of Veterinary Pathology, Krantisinh Nana Patil College of Veterinary Science, Shirwal, Maharashtra, Dr. Dilip Patil, Scientist D, ICMR-NIV, Pune and the laboratory team of Maximum Containment Facility, ICMR-NIV, Pune which includes Mr. Shreekant Baradkar, Mr. Annasaheb Suryawanshi, Mr. Rajen Lakra, Mrs. Ashwini Waghmare, Ms. Manisha Dudhmal. Mr. Kundan Wakchuare, Mr Sanjay Gopale, Ms. Tejashri Kore, Ms. Shilpa Ray, Ms. Priyanka Waghmare, Ms. Poonam Bodke, Mr. Madhav Acharya, Mr. Sanjay Thorat and Mr. Ganesh Chopade.

## Author contributions

**Conceptualization:** PDY, SM, SK

**Methodology:** SM, PDY, GD, GS, ASA

**Investigation:** PDY, SM, GS, ASA, DN, MK, AK, PS, PG

**Visualization:** PDY, SM, SK, GS, ASA, DYP

**Supervision:** PDY, SM, GS, ASA

**Writing - original draft:** PDY, SM

**Writing – review & editing:** PDY, SM, GS, ASA, GD, DN, SK, DYP, PA

## Data and materials availability

All data are available in the main text or the supplementary materials.

